# Cryo-electron microscopy structures of pyrene-labeled ADP-P_i_- and ADP-actin filaments

**DOI:** 10.1101/2020.08.26.268748

**Authors:** Steven Z. Chou, Thomas D. Pollard

## Abstract

We report high resolution cryo-electron microscopy structures of actin filaments with N-1-pyrene conjugated to cysteine 374 and either ADP (3.2 Å) or ADP-phosphate (3.0 Å) in the active site. Polymerization buries pyrene in a hydrophobic cavity between subunits along the long-pitch helix with only minor differences in conformation compared with native actin filaments. These structures explain how polymerization increases the fluorescence 20-fold, how myosin and cofilin binding to filaments reduces the fluorescence and how profilin binding to actin monomers increases the fluorescence.

## Introduction

For almost forty years, N-(1-pyrene) iodoacetamide has been used to label actin at C374, the only solvent-accessible of five cysteine residues (Kouyama and Mihashi 1981). Hundreds of studies have used the fluorescence of pyrenyl-actin to measure actin polymerization (Kouyama and Mihashi 1981, Cooper, Walker et al. 1983) and binding of myosin (Kouyama and Mihashi 1981, Taylor 1991), profilin (Lee, Li et al. 1988) and cofilin (Nishida, Maekawa et al. 1984, Carlier, Laurent et al. 1997) to actin filaments and monomers. The mechanisms of the fluorescence changes are still unknown due to the lack of structural information.

Here we provide high-resolution cryo-electron microscopy structures of pyrene-labeled ADP-P_i_- and ADP-actin filaments, which not only explain these mechanisms but also show how the presence of pyrene modifies the filament structure in small ways and why other dyes coupled to C374 compromise polymerization. We also report that phosphate binding increases the fluorescence of to pyrenyl-Mg-ADP-actin filaments.

## Results

### Cryo-EM structures of pyrenyl-actin filaments

Image processing of electron micrographs of 411,301 particles of Mg-ADP-P_i_-actin filaments produced an electron potential map with a resolution 3.0 Å and 240,254 particles of Mg-ADP-pyrenyl-actin filaments gave a map with a resolution of 3.2 Å (Fig. S2; Table S1). These maps showed the positions of most side chains, the conjugated pyrene, bound nucleotides and associated Mg^2+^ and allowed unambiguous model building (Fig. 1). The helical parameters (rise and twist) of pyrenyl-actin filaments are nearly identical to native actin filaments in both the ADP-P_i_ and ADP states (Table S2). The conformations of the bound nucleotides and their surrounding residues are the same in pyrenyl-actin filaments and native actin filaments (Fig. 1C & D).

**Figure 1.**
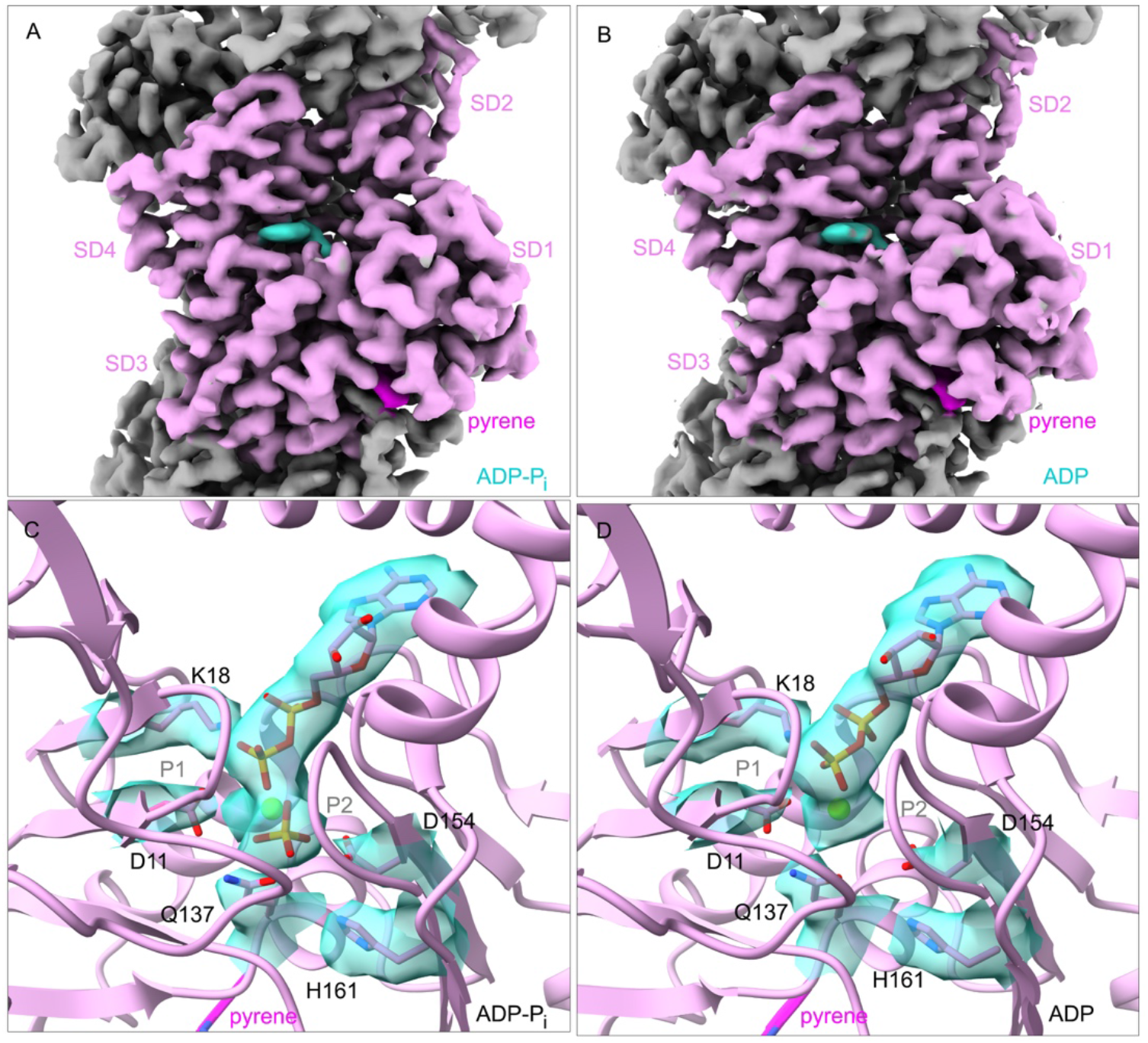
Electron potential maps and ribbon diagrams of pyrenyl-actin filaments with (A, C) bound Mg-ADP-P_i_ or (B, D) bound Mg-ADP. (A-B) Maps with highlighted densities of the nucleotide (turquoise) and pyrene (magenta) in one subunit (plum). Neighboring subunits are grey. The barbed end is at the bottom. The four subdomains are labeled as SD1-4. (C-D) Ribbon models with the densities and stick figures of the bound nucleotides and a few key neighboring side chains. P1 (residues 11-16) and P2 (residues 154-161) are the two phosphate binding loops.

The maps of both ADP-P_i_- and ADP-pyrenyl actin filaments show a clear, continuous extra density for the conjugated pyrene group (Fig. 2 A & B). The pyrene group conjugated to C374 is largely sandwiched in a fixed conformation between two adjacent actin subunits along the long-pitch helix between the barbed end groove of the subunit near the pointed end (called the P subunit hereafter) and D-loop of the subunit near the barbed end (called the B subunit hereafter). Large unidentified densities also appeared in this location in maps of actin filaments with bound ADP-BeF_x_ and bound jasplakinolide with either ADP-P_i_ or ADP published by Merino et al. (Merino, Pospich et al. 2018).

**Figure 2.**
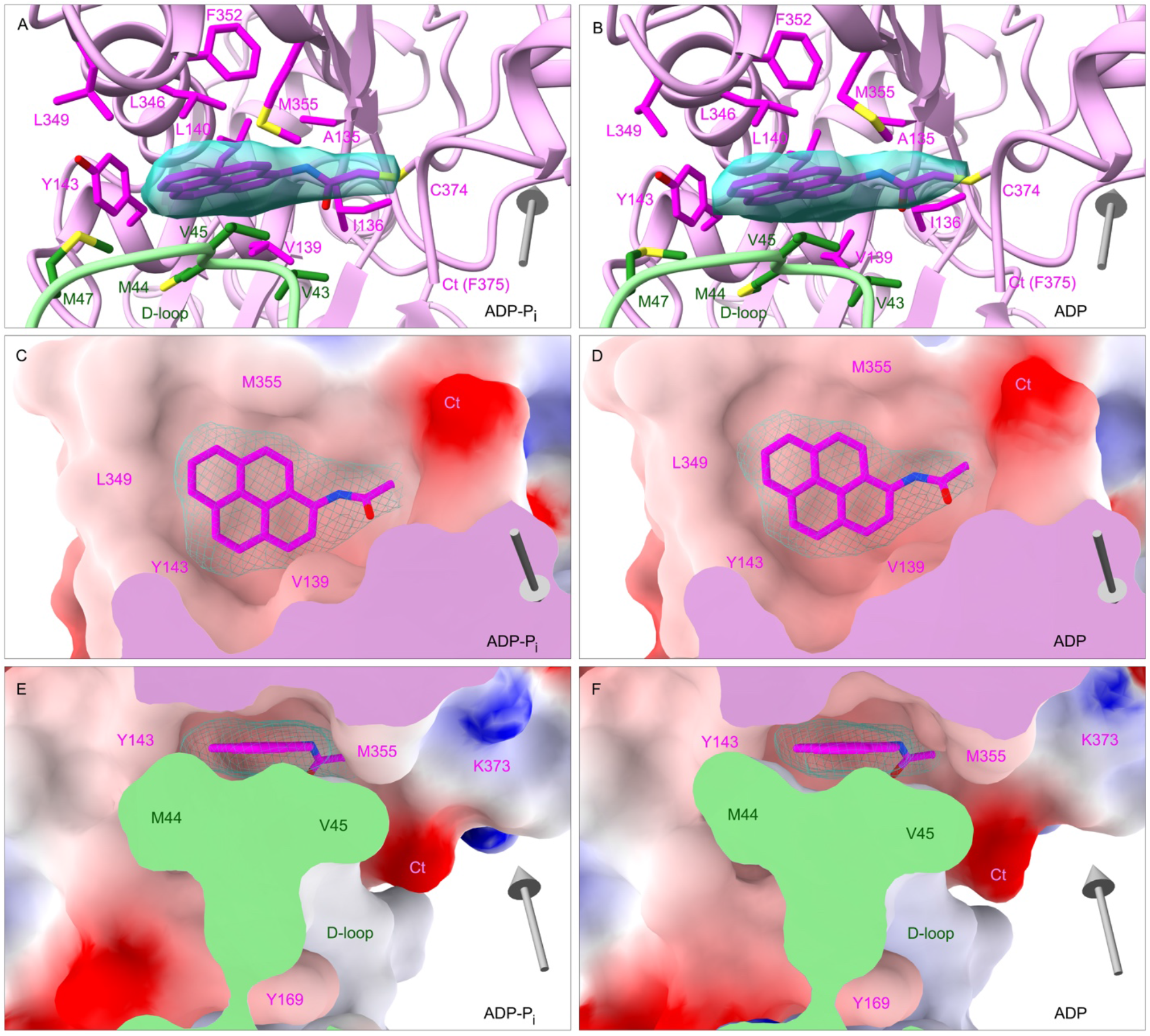
N-(1-pyrene) conjugated to C374 in a hydrophobic pocket between two subunits along the long-pitch helix of the actin filament. EM map densities of N-(1-pyrene) are shown as turquoise mesh surfaces and N-(1-pyrene) as magenta stick figures. Grey arrows show the filament orientation with arrow heads pointing to the pointed end. (A, C, E) filaments with bound Mg-ADP-P_i_; (B, D, F) filaments with bound Mg-ADP. (A, B) Backbone ribbon diagrams. The actin subunit toward the pointed end is colored plum and the subunit toward the barbed end is light green. (C, D, E, F) Electrostatic potential surfaces (blue: positive; red: negative) of interacting actin subunits. (C, D) Face views (E, F) and side views. Neighboring subunits in C and D are omitted for clarity. The cut surfaces of the two actin subunits in the section views are colored in plum and light green.

The pyrene group in the filaments sits snugly in a hydrophobic pocket (Fig. 2C-F). Electrostatic potential maps calculated from the refined models show that the pyrene group is surrounded by the hydrophobic residues A135, I136, V139, L140, L346, L349, F352, and M355 in the barbed end groove of the P subunit and V43, M44, V45, M47 of the D-loop of the B subunit. This environment differs from the actin monomer where the pyrene group of C374 is exposed, at least in part, to the water solvent. This change of environment likely explains why the fluorescence intensity of pyrenyl actin increases ∼20 folds upon polymerization.

### Structural difference between pyrenyl and native actin filaments

Superimposition of the maps and models of the pyrenyl-actin filaments and native actin filaments showed that the presence of the pyrene group has a larger impact on longitudinal than lateral interactions between the subunits (Fig. 3). Pyrene displaces by small amounts the side chains of Y143 and V45 and the backbones of residues 44-48 of the D-loop of the B subunit, leaving residues 40-43 and 49-50 at either end of the D-loop in place (Fig. 3F). The pyrene group attached to the side chain of C374 also changes the local conformations of the C-terminal residues 373-375. The only difference in lateral interactions between subunits in the presence of pyrene was the movement of the side chain of K113 from about 5.0 Å to its own C-terminal carboxyl group to about 11.0 Å.

**Figure 3.**
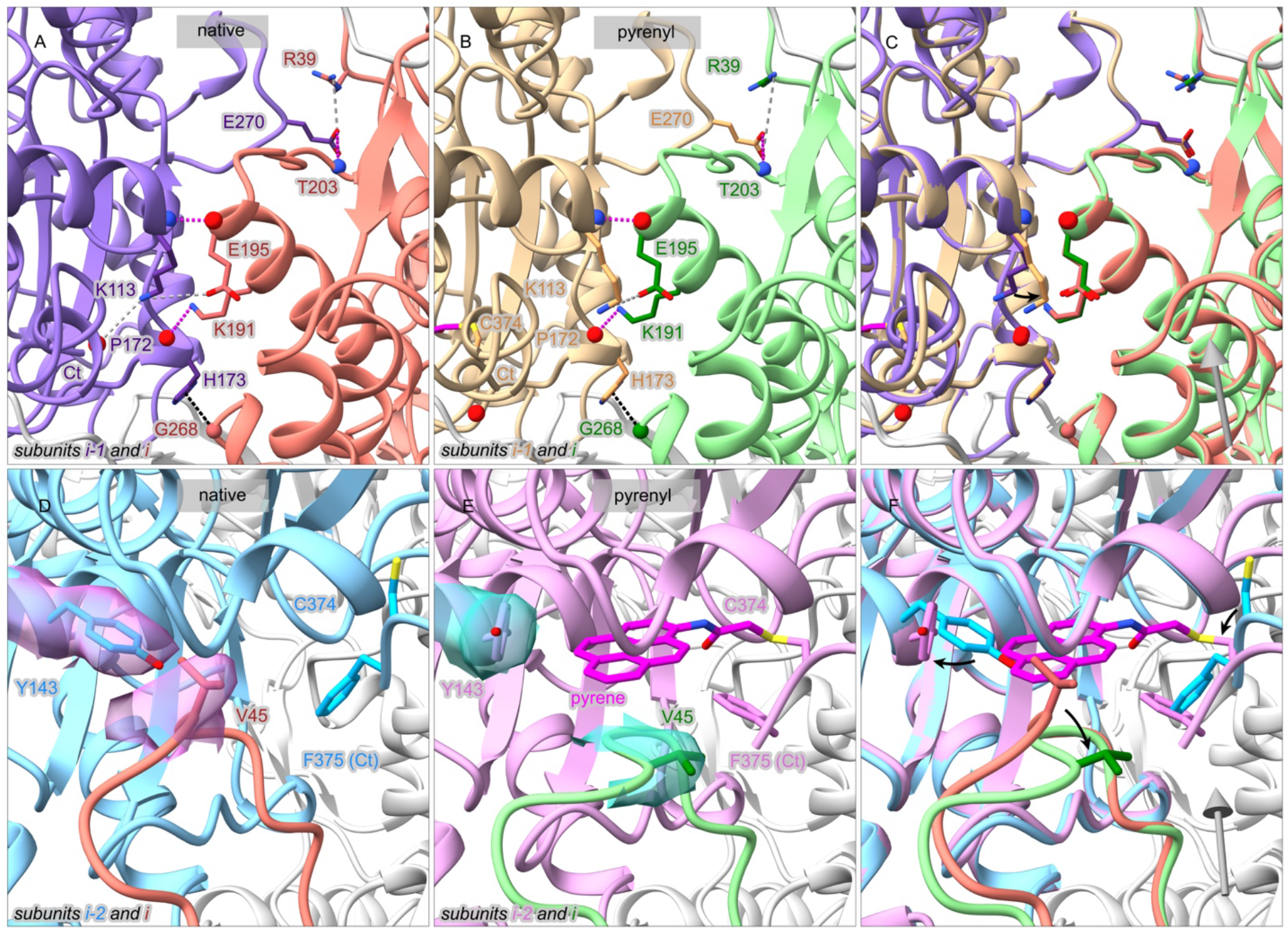
Influence of pyrene labeling of C374 on subunit interactions in actin filaments. Ribbon diagrams have stick figures of important side chains and balls for important backbone nitrogen atoms (blue), oxygen atoms (red) and C_α_ atoms (green or salmon). Grey arrows on the right column have their arrow heads pointing along the filament axis toward the pointed end. Subunits in native actin filaments are colored sky blue (subunit *i-2*), medium purple (subunit *i-1*), and salmon (subunit *i*); subunits in pyrenyl-actin filaments are colored plum (subunit *i-2*), tan (subunit *i-1*), and light green (subunit *i*). Dashed lines show charge-charge interactions (grey), hydrogen bonds (magenta) and *van der Waals* contacts (black). (A, B, C) Pyrene does not change Most lateral interactions along the short-pitch helix are the same without and with pyrene, but in pyrenyl-actin the side chain of K113 rotates (black arrow) and loses an electrostatic interaction with its own C-terminal carboxyl group while retaining an electrostatic interaction with the side chain of E195 of the neighboring subunit. (D, E, F) Effects of pyrene labeling on interactions along the long-pitch helix. Black arrows mark how pyrene causes localized changes of the conformations of V45 in the D-loop and the side chain of Y143 in the adjacent subunit. Insertion of pyrene between two actin subunits displaces the C_α_ atom of V45 towards the filament barbed end by 2.9 Å and rotation of the side chain 44.1° without disturbing other interactions between D-loop and the neighboring subunit. Pyrene twists the aromatic ring of Y143 about 72.3°.

### Effect of phosphate in the nucleotide binding site of ADP-actin filaments on labeling with N-(1-pyrene) iodoacetamide and the fluorescence intensity of pyrenyl-actin filaments

The fluorescence emission of N-(1-pyrene) iodoacetamide is higher after conjugating to actin filaments (Fig. 4A). The conjugation reaction is very slow with half times >2 h, indicating that C374 is poorly accessible. The solvent accessible surface area of the side chain of C374 is ∼0.1 Å^2^ in native ADP-P_i_-actin filaments and ∼6.2 Å^2^ in native ADP-actin filaments (Chou and Pollard 2019). Labeling was slower by a factor of 2 in the presence of phosphate than with sulfate or no additions.

**Figure 4.**
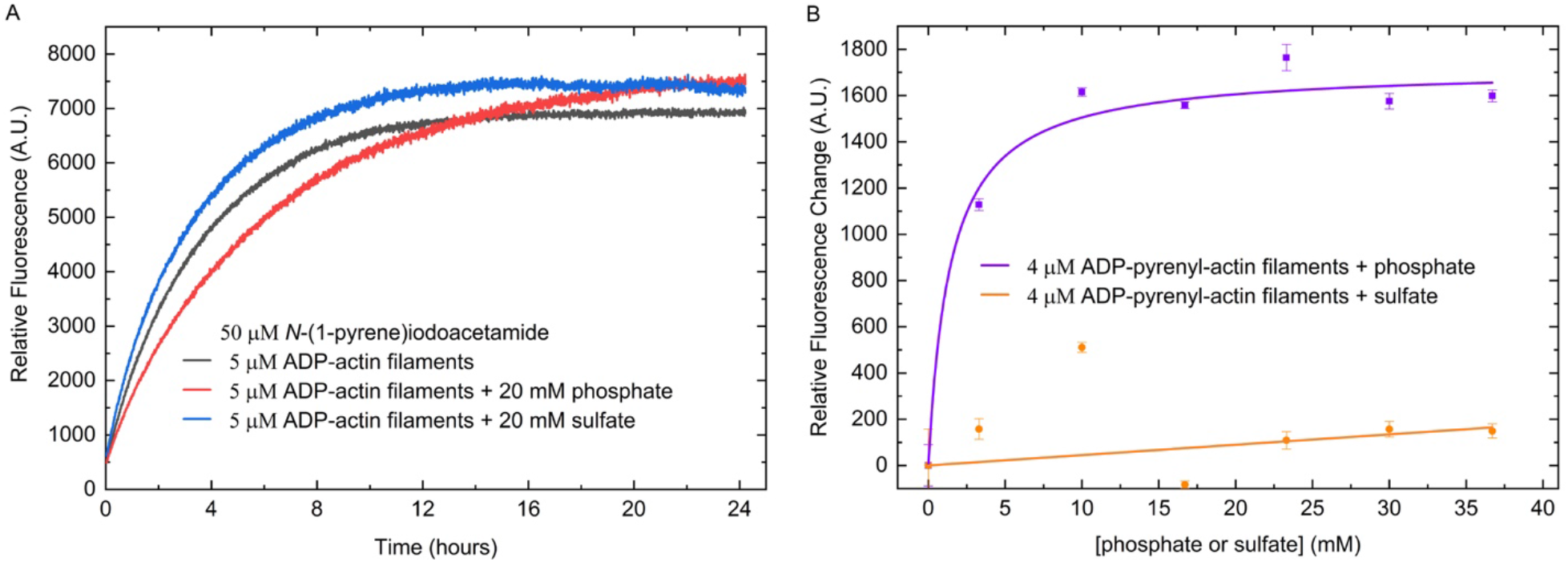
Interactions between phosphate in the nucleotide binding site and pyrene. (A) Phosphate slows the labeling of ADP-actin filaments with N-(1-pyrene)iodoacetamide. Solutions of 5 µM Mg-ADP-actin filaments were polymerized overnight at 4°C in 100 mM KCl; 1 mM MgCl_2_; 10 mM imidazole, pH 7.0; 0.3 mM ADP; 3 mM NaN_3_ and preincubated at room temperature with either no additions, 20 mM potassium phosphate or 20 mM potassium sulfate before adding 50 µM N-(1-pyrene)iodoacetamide. The basal fluorescence is from the free N-(1-pyrene)iodoacetamide, and the fluorescence increase is due to conjugation of N-(1-pyrene)iodoacetamide to the sidechain of C374 in actin filaments. (B) Effect of phosphate in the buffer on the fluorescence of Mg-ADP-pyrenyl-actin filaments. Each data point was acquired on a sample consisting of 120 µL of 5 µM Mg-ADP-pyrenyl-actin filaments and 30 µL of phosphate or sulfate at a specific concentration. Each sample was incubated for ∼1 hour before the measurements. The Y-axes are in arbitrary units (A.U.).

The fluorescence intensity of Mg-ADP-pyrenyl-actin filaments increases with the concentration of phosphate in the buffer (Fig. 4B) corresponding to a K_d_ <10 mM, similar to other measurement of the affinity of the filaments for phosphate (Carlier and Pantaloni 1986).

### Differences near the pyrene group in Mg-ADP-P_i_- and Mg-ADP-pyrenyl-actin filaments

The RMSDs between the alpha-carbons of the models for Mg-ADP-P_i_- and Mg-ADP-pyrenyl-actin filaments were only 0.35 Å, smaller than the RMSDs of native actin filaments with the two nucleotides, but comparison of the maps revealed subtle differences at the C-terminus and D-loop (Fig. 5). The densities were weaker in Mg-ADP-pyrenyl-actin filaments than Mg-ADP-P_i_-pyrenyl-actin filaments for the connection between the side chain of C374 and pyrene, F375 and the backbone of G48 (Fig. 5A & C). These missing or weak densities indicate that the longitudinal interface between the B and P subunits is less well ordered in Mg-ADP-pyrenyl-actin filaments than Mg-ADP-P_i_-pyrenyl-actin filaments.

**Figure 5.**
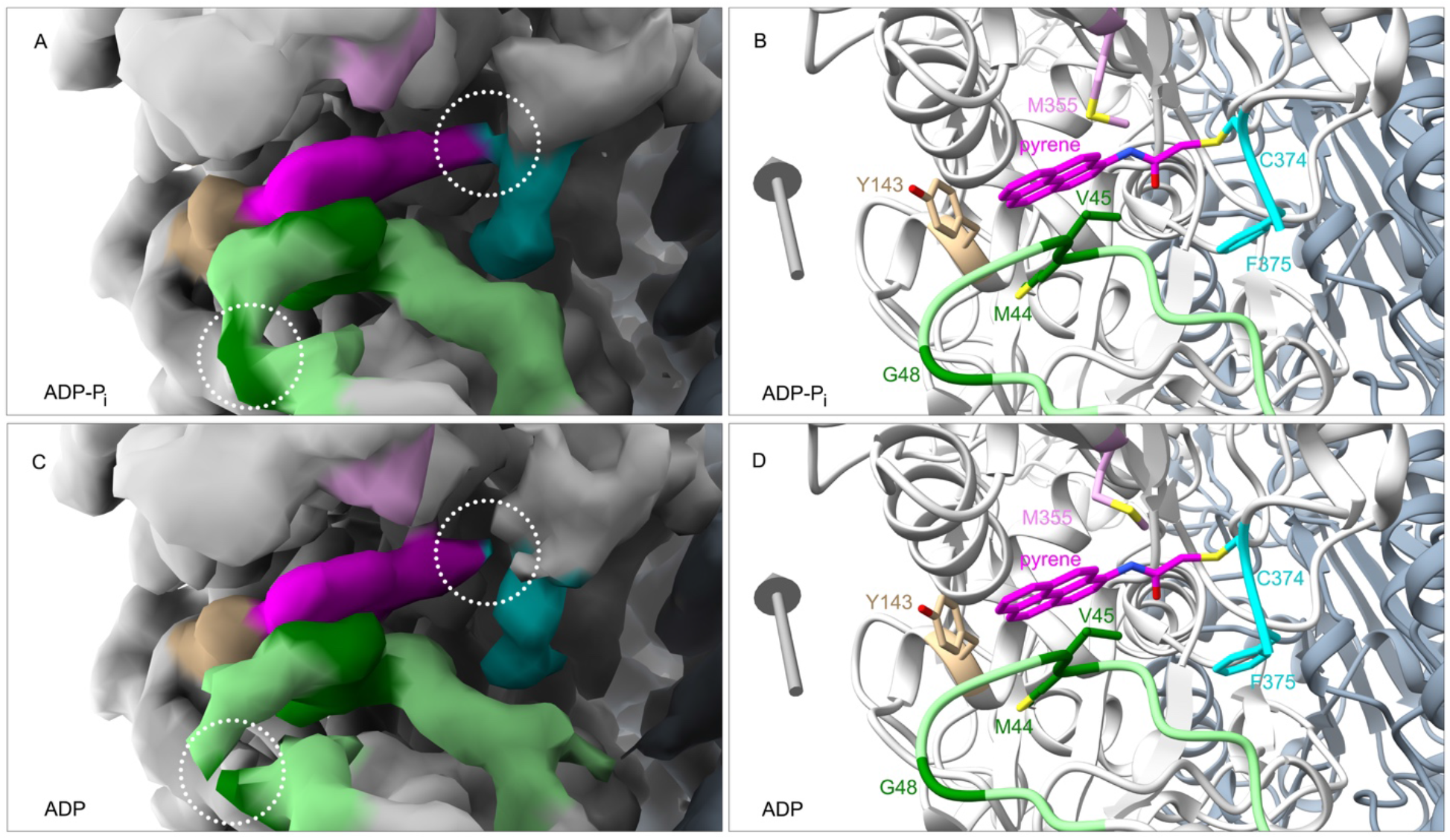
Subtle differences in the pyrene and surrounding residues in (A-B) Mg-ADP-P_i_- and (C-D) Mg-ADP-pyrenyl-actin filaments with map densities on the left and ribbon diagrams on the right. Both maps are contoured at the same level (0.015 V). Color code: actin subunits, light grey in one strand and slate grey in the other strand; D-loop (residues 40-50) of the adjacent actin subunit, light green with residues M44, V45 and G48, dark green; pyrene, magenta; C-terminus including C374 and F375, cyan; Y143, tan; M355, plum. Grey arrows are orientated with their heads pointing to the pointed end of the filaments. In the ADP-actin filament (C), note the lower/broken density for G48 and the connection between pyrene and the side chains of C374.

### Interactions of myosin and cofilin with pyrenyl-actin filaments and profilin with pyrenyl-actin monomers

The original application of pyrenyl-actin filaments was to measure myosin binding (Kouyama and Mihashi 1981), because bound myosin reduces the fluorescence intensity by ∼30%. The presence of pyrene also reduces the affinity of myosin for actin filaments by about half (Taylor 1991). Superimposing the structure of a native ADP-actin filament decorated with myosin-Ib (PDB ID: 6C1D) onto our ADP-pyrenyl-actin filament showed that the presence of pyrene forces the side chain of actin Y143 into a position that slightly clashes with the side chain of myosin P467 and the backbone of actin M47 into a minor clash with the side chains of myosin R466 and K478 (Fig. 6B). Reciprocally, when myosin is bound to actin, the region of the actin D-loop around V45 would clash with the pyrene (Fig. 6B). Therefore, strong binding of myosin to actin filaments likely forces Y143 and the D-loop to displace the pyrene partially from its hydrophobic pocket into the solvent, accounting for the lower fluorescence.

**Figure 6.**
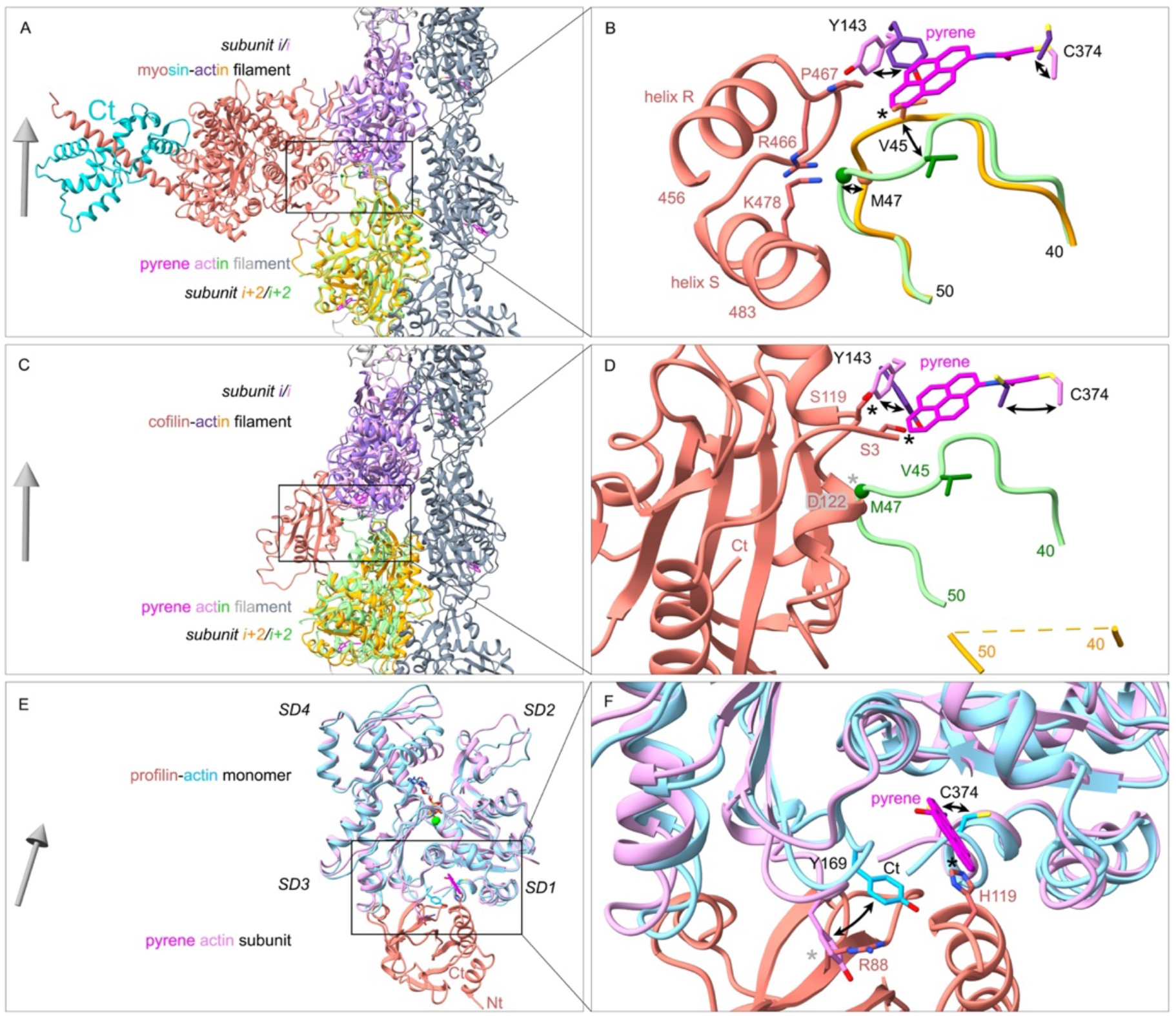
Ribbon diagrams showing the influence of pyrene labeling on the interactions of myosin and cofilin with actin filaments and of profilin with actin monomers. Grey arrows show the filament orientation with arrowheads pointing to the pointed end. Black curved arrows show conformational changes. Black stars label strong clashes; grey stars mark local conformational changes that cause minor or no clashes. (A, C, E) overviews; (B, D, F) simplified close-up views. (A, B) Superimposition of an ADP-actin filament decorated with myosin-Ib (PDB ID: 6C1D) on our pyrenyl-ADP-actin filament. All the C_α_ atoms of subunit *i* in these two filaments were used for the superimposition. For clarity, only two actin subunits (*i* and *i+2*) in myosin-decorated ADP-actin filament are displayed. The presence of pyrene would cause two steric clashes with myosin: rotation of the side chain of Y143 of actin would clash with the cyclic side chain of P467 of the myosin heavy chain; and the backbone of actin M47 clashes with the side chains of myosin R466 and K478. Binding of myosin to pyrenyl actin filament is likely to displace pyrene partially from its hydrophobic pocket. (C, D) Superimposition of a cofilin-decorated ADP-actin filament (PDB ID: 5YU8) on our pyrenyl-ADP-actin filament. The C_α_ atoms of residues in subdomain 3 of subunit *i* were used for the superimposition. The presence of pyrene would cause two steric clashes with cofilin: the side chain of actin Y143 clashes with the side chain of cofilin S119; and the pyrene group clashes with the side chain of cofilin S3. Binding of cofilin to pyrenyl ADP-actin filament would squeeze part of the pyrene group out of the hydrophobic pocket and displace the D-loop of the neighboring subunit. (E, F) Superimposition of a profilin-actin monomer complex (PDB ID: 2BTF) on a subunit in our pyrenyl-ADP-actin filament using subdomains 1 and 3. The C_*α*_ atoms of residues in subdomains 1 and 3 are used for their superimposition. The side chain of profilin H119 would clash with the pyrene group. The side chain of actin Y169 could avoid the clash with the side chain of R88 in profilin by rearranging the conformation of the Y169 loop (WH2-binding loop).

Cofilin binding reduces the fluorescence of Mg-ADP-pyrenyl-actin filaments more than ten-fold without depolymerization (Nishida, Maekawa et al. 1984, Carlier, Laurent et al. 1997). Superimposing the structure of one subunit from a cofilin-decorated native ADP-actin filament (PDB ID: 5YU8) onto our ADP-pyrenyl-actin filament showed that the presence of pyrene would cause two strong steric clashes with cofilin: the side chain of actin Y143 clashes with the side chain of cofilin S119; and the pyrene group clashes with the side chain of cofilin S3 (Fig. 6D). To avoid this steric clash, cofilin must push Y143 toward the pyrene, which would displace pyrene from the hydrophobic pocket (Fig. 6D). This exposure of the pyrene group in the cofilin-decorated ADP-pyrenyl-actin filament would likely reduce its quantum yield. Furthermore, most of the D-loop residues (41-49) are disordered in cofilin-decorated ADP-actin filaments, eliminating the bottom of the hydrophobic pocket for pyrene.

Profilin binding enhances the fluorescence of pyrenyl-actin monomers (Lee, Li et al. 1988), but the pyrene label reduces the affinity of actin monomers for profilin by an order of magnitude (Vinson, De La Cruz et al. 1998). Superimposing a crystal structure of profilin bound to native ATP-actin monomer (PDB ID: 2BTF) onto a subunit from our ADP-pyrenyl-actin filament showed that the side chain of H119 in profilin would clash with the pyrene group, accounting for the lower affinity of profilin for pyrenyl-actin monomers. The side chain of actin Y169 could avoid the clash with the side chain of profilin R88 by rearranging the conformation of the Y169 loop (WH2-binding loop). This increases the buried surface area of pyrene from 295 Å^2^ in the actin monomer to 356 Å^2^ in profilin-actin monomer. This together with a more hydrophobic environment for the pyrene may account for the modestly higher fluorescence.

## Discussion

### Mechanism of fluorescence enhancement by polymerization of pyrenyl-actin monomers

This change of environment from a solvent-exposed position to a hydrophobic pocket between the subunits in a filament explains why the fluorescence intensity of pyrenyl actin increases ∼20 folds upon polymerization. Pyrene fits almost perfectly into this hydrophobic pocket but displaces the side chain and backbone of V45. The side chain of M44 is the most important residue for inter-subunit interactions along the long-pitch helix. The presence of the pyrene group moves the backbone C_α_ atom of M44 toward the filament barbed end by 0.7 Å, but its side chain remains in place. These small changes do not alter the critical concentration or kinetics of spontaneous polymerization.

On the other hand, labeling actin residue C374 with tetramethylrhodamine or Oregon green (Fig. S4) alters the rates of Mg-ATP-actin monomers binding and dissociating at the ends of actin filaments with the changes depending on the fraction of labeled subunits (Amann and Pollard 2001, Fujiwara, Zweifel et al. 2018). Both of these fluorescent dyes are much larger than pyrene, so they cannot fit into the hydrophobic pocket between subunits along the long-pitch helix and are likely to clash with the adjacent subunit. The positive charge on tetramethylrhodamine may contributing to inhibiting elongation more strongly than Oregon green. Labeling lysine with a fluorescent dye has smaller effects on polymerization (Fujiwara, Zweifel et al. 2018).

### Mechanisms of fluorescence quenching by myosin and cofilin

Binding of both myosin and cofilin to pyrenyl-actin filaments involves steric clashes that do not occur in native actin filaments (Fig. 6B & D). These clashes likely result in local conformational changes that expose the pyrene to solvent and explain why both proteins lower the fluorescence and why the presence of pyrene reduces the affinities of both proteins for actin filaments. Strong binding of myosin to actin filaments likely force Y143 and the D-loop to displace the pyrene partially from its hydrophobic pocket into the solvent, accounting for the lower fluorescence. To bind an actin filament, cofilin must push actin Y143 toward the pyrene, which would partially displace pyrene from the hydrophobic pocket.

Profilin binding enhances the fluorescence of pyrenyl-actin monomers. The likely explanation is reducing the exposure of pyrene to the solvent. Minor steric clashes between profilin and actin labeled with pyrene explain why the dye reduces the affinity ten-fold.

### Effect of phosphate on the labeling and fluorescence of ADP-actin filaments

Phosphate associated with ADP in the active site influences both the reaction of N-(1-pyrene) iodoacetamide with actin filaments and the fluorescence of pyrenyl-actin filaments. In our high-resolution EM structures of native actin filaments (Chou and Pollard 2019), C374 is more exposed in the ADP-actin filament than the ADP-P_i_-filament, explaining the different labeling rates. The explanation for the effect of phosphate on the fluorescence of pyrenyl-ADP-actin filaments is less clear, but may be due to the pyrene being trapped more rigidly in the hydrophobic pocket of filaments with phosphate. Lower map densities for the link between C374 and pyrene and around G48 are evidence of greater mobility in the ADP-actin filaments.

### Chemical and post-translational modifications of C374

Actin C374 can be post-translationally modified by oxidation, glutathionylation, carbonylation and nitrosylation (Terman and Kashina 2013). These modifications on C374 usually decrease the polymerization rate, increase the critical concentration, weaken filament stability and alter actin functions. These modifying chemical groups, especially large hydrophobic groups like glutathione, may occupy the space where pyrene binds.

## Methods and Materials

### Materials

We purchased N-(1-pyrene)iodoacetamide (sc-472510) from Santa Cruz Biotechnologies, ATP (A2383), ADP (A2754), hexokinase (H6380) from Sigma-Aldrich, and glucose (167454) from Thermo Fisher Scientific.

### Actin Purification

Chloroform-washed actin acetone powder was made from the flash-frozen chicken skeletal muscle (Feuer and Straub 1948) purchased from a local Trader Joe’s grocery store. Actin was extracted from the powder with buffer G (2 mM Tris-HCl, pH 8.0, 0.1 mM CaCl_2_, 0.1 mM ATP, 0.5 mM DTT, 1 mM NaN_3_) for 30 min at 4°C, and polymerized by adding KCl to 50 mM and MgCl_2_ to 2 mM. Actin filaments were pelleted by centrifugation at 140,000*× g* for 2 hours at 4°C and depolymerized by dialysis against four changes of buffer G at 4°C over 72 h. The sample was centrifuged at 140,000*× g* for 2 hours at 4°C. The top 2/3 of depolymerized actin supernatant was purified by gel filtration on a Sephacryl S-300 column equilibrated with buffer G (MacLean-Fletcher and Pollard 1980). The concentration of native actin monomers was determined by absorbance at 290 nm using [actin, µM] = A_290_ *×* 38.5.

### Labeling ADP-actin filaments with N-(1-pyrene) iodoacetamide

The bottom 1/3 of depolymerized actin supernatant was polymerized by dialyzing for 10 h against three changes of PL buffer (100 mM KCl, 2 mM MgSO_4_, 25 mM Tris-HCl, pH 7.5, 0.3 mM ATP, 3 mM NaN_3_). During polymerization, bound ATP was hydrolyzed into ADP and phosphate (P_i_). Dissociated P_i_ and DTT (from buffer G) were removed during dialysis. The repolymerized actin was diluted to 1 mg/mL (23.8 µM) with PL buffer and incubated at 4°C for 36 h with 0.15 mM N-(1-pyrene)iodoacetamide from a stock solution of 10 mM dissolved in dimethylformamide. Labeled actin filaments were collected by centrifugation, the pellet was homogenized in buffer G, and the mixture dialyzed against buffer G for 2 days at 4°C. After clarification at 4°C in a Beckman Ti70 rotor at 38,000 rpm for 2 h, the top 80% of supernatant was gel filtered through a Superdex 200 column equilibrated with buffer G. The concentrations of pyrene and actin in the peak tail fractions were determined by measuring the absorbances at 344 nm (A_344_) and 290 nm (A_290_) and using these formulas: [pyrene, µM] = A_344_ *×* 45.5 µM/OD and [actin, µM] = (A_290_ – (A_344_ *×* 0.127)) *×* 38.5 µM/OD (Kouyama and Mihashi 1981, Cooper, Walker et al. 1983). The ratio of pyrene/actin was close to 1.0. Both native and pyrenyl actin monomers were stored at 4°C and used in less than a week.

Ca^2+^ bound to actin monomers was exchanged for Mg^2+^ by incubating with 0.1 volumes of 10*×* ME buffer (0.5 mM MgCl_2_, 2 mM EGTA, pH 7.5) at room temperature for 10 min. Mg-ATP-pyrenyl-actin was converted to Mg-ADP-pyrenyl-actin by incubating with 1 mM glucose and 5 units/mL hexokinase at room temperature for 30 min, followed by adding 1 mM ADP, pH 7.0 and polymerizing the actin for 2 h at 25°C by adding 0.1 volumes of 10*×* KMEI buffer (1 M KCl, 10 mM MgCl_2_, 10 mM EGTA, 100 mM imidazole, pH 7.0, 5 mM DTT) to make ADP-actin filaments or first adding potassium phosphate (pH 7.0) to 30 mM and then 0.1 volumes of 10*×* KMEI buffer to make ADP-P_i_-actin filaments. Pyrenyl actin filaments were stored at 4°C for 10 h before vitrification for EM analysis.

### Fluorescence assay to measure pyrene labeling

Mg-ADP-actin filaments (94 µL; 10.7 µM) in PLD buffer (100 mM KCl, 2 mM MgCl_2_, 10 mM imidazole, pH 7.0, 0.3 mM ADP, 3 mM NaN_3_) were preincubated by adding 6 µL of either H_2_O, 250 mM potassium phosphate pH 7.0, or 250 mM K_2_SO_4_ pH 7.0 for 10 min before adding 50 µL of 150 µM N-(1-pyrene)iodoacetamide diluted from the above-mentioned stock solution with PLD buffer. The time course of the fluorescence change was measured in a Costar black polystyrene 96-well plate (3694; Corning Incorporated, Corning, NY) using a SpectraMAX plate reader (Genimi EM; Molecular Devices, San Jose, CA) with excitation at 365 nm and emission at 407 nm.

### Sample Freezing and Image Collection

Holy carbon Quantifoil 2/1 300-mesh gold grids (Electron Microscopy Sciences, Hatfield, PA) were glow-discharged for 30 s in a Bal-Tec SCD 005 sputter coater (Leica Biosystem Inc.) in air (pressure: 0.05 mBar) at 25 mA. We applied 3 µL of 18 µM of pyrenyl actin filaments to the carbon side of the grids in a Vitrobot Mark IV (FEI company, Hillsboro, OR) at 10°C and 100% humidity. After waiting 40 s, extra solution was blotted off using Vitrobot standard filter paper (grade: 595; Ted Pella, Redding, CA) for 2.5 s at blot force −15. Grids were frozen by plunging into liquid ethane cooled to about −180°C and screened on a Talos electron microscope operated at 120 kV and equipped with a Ceta 16M camera (FEI company, Hillsboro, OR). The two datasets were collected in a consecutive session on a Titan Krios microscope equipped with an XFEG at 300 kV, a nanoprobe, and a Gatan image filter (slit width: 20 eV). Movies were recorded at a series of defocus values between −1.0 µm and −2.5 µm on a K2 summit camera in super-resolution mode, using the beam image shift strategy (4 movies/hole) designed in the third-party program SerialEM (Mastronarde 2005). Each movie had 55 frames and each frame time was 0.25 s. The dose rate on camera level was 5.6 counts/pixel/s, and the physical pixel size was 1.045 Å.

We collected 1560 low-dose movies of Mg-ADP-P_i_-actin filaments and 1980 low-dose movies of Mg-ADP-pyrenyl-actin filaments. The ADP-actin filament samples (critical concentration: 0.8 µM) had more actin monomers in the background than the ADP-P_i_-actin filament sample (critical concentration 0.1 µM) (Fig. S1 A & B).

### EM Map Reconstruction

Movies were dose-averaged, magnification-corrected, motion-removed, and summed with MotionCor2 (Zheng, Palovcak et al. 2017) using 9 *×* 9 patches. Parameters in the contrast transfer function (CTF) were estimated with Gctf (Zhang 2016) using summed but unweighted micrographs. Filaments were first autopicked in RELION (He and Scheres 2017) with a curve factor of 0.95. The coordinates were exported in box format for manual adjustment with sxhelixboxer.py in SPARX (Hohn, Tang et al. 2007). The manually-adjusted filament coordinates were read back into RELION for further analysis. Before helical reconstruction, the filaments were windowed into overlapping square segments by translating the box (328 *×* 328 pixels) along the central axis of each filament by one subunit (26 pixels). Each segment contained ∼12 subunits. Our map of native ADP-P_i_-actin filaments (EMD-7937) was low-pass filtered at 10 Å before being used as the initial map. Particle local CTFs were used in the reconstructions. A soft-edged 3D mask with a radius of 45% of the box size was created for postprocessing. The B-factors for map sharpening were determined by RELION. A cutoff of 0.143 was used for resolution estimation using the Fourier shell correlation method. At this point, the map of Mg-ADP-P_i_-pyrenyl-actin filaments was refined to 3.3 Å, and that of Mg-ADP-pyrenyl-actin filaments was 3.5 Å.

Because the filaments are continuous, the particle local CTFs should change continuously along each filament. However, the particle local CTFs varied along most of the filaments owing to the low signal-to-noise ratio in the images, a nature of electron micrographs. Therefore, we smoothed the CTF of each particle along the same filament locally over five neighboring particles. The highest and lowest CTFs were removed and the remaining three were fitted using linear regression. After this smoothing step, most of particle local CTFs changed continuously along each filament.

We calculated the rise and twist for the consistently aligned particles (>99.8%), and then calculated their mean and standard deviation. After removing the rise and twist outliers (>2*×* SD), we reconstructed a new map using particles in a certain confidence interval (95%). The best reconstructions were obtained after a trade-off between particle number and assembly homogeneity. This process improved the resolutions of the maps to 3.0 Å for Mg-ADP-P_i_-pyrenyl-actin filaments and to 3.2 Å for Mg-ADP-pyrenyl-actin filaments (Fig. S2; Table S1). Local resolutions were estimated with ResMap (Kucukelbir, Sigworth et al. 2014). All the calculations were run on the Farnam computer cluster maintained by Yale High Performance Computing team.

### Model Building, Refinement and Visualization

Our EM structure of native ADP-P_i_-actin filaments (PDB ID: 6DJN) was used as the initial model. The chemical restraints for the pyrene moiety in cif format were generated with eLBOW tool in Phenix (Moriarty, Grosse-Kunstleve et al. 2009, Adams, Afonine et al. 2010). The atomic coordinates for the pyrene moiety were named according to the crystal structure of N-(1-pyrene)acetamide-labeled P450 (PDB ID: 4KKY). The coordinates of actin and pyrene were joined together manually in a text editor, and the pyrene group was moved close to C374 of each actin subunit in Coot (Emsley, Lohkamp et al. 2010) to get the starting model of pyrenyl-actin filaments. Further restraints for linking the pyrene moiety to actin in edits format were added manually. The coordinates were refined against the EM maps of pyrenyl-actin filaments in real space with Phenix (Afonine, Poon et al. 2018) and checked with Coot. Figures of maps and models were generated with ChimeraX (Goddard, Huang et al. 2018). The electrostatic potentials were calculated with DelPhi (Rocchia, Sridharan et al. 2002), and mapped onto the molecular surfaces in ChimeraX using an offset of 0 Å. Root mean square deviations and buried surface areas were calculated with Chimera (Pettersen, Goddard et al. 2004), and solvent accessible surface areas were calculated with PyMOL (Schrodinger 2015).

## Acknowledgements

Research reported in this publication was supported by National Institute of General Medical Sciences of the National Institutes of Health under award number R01GM026338. The content is solely the responsibility of the authors and does not necessarily represent the official views of the National Institutes of Health. The authors thank Drs. Shenping Wu, Marc Llaguno, Xinran Liu Kaifeng Zhou and Paul Raccuia for managing the electron microscopes, Ed Taylor for his ideas and suggestions, Mark Mooseker for using his equipment and Shoshana Zhang and Aharon Rosenbloom help with fluorescence experiments. The authors also thank the Yale Center for Research Computing for guidance and the use of the research computing infrastructure, specifically the Farnam cluster.

## Author contributions

S.Z.C. and T.D.P. designed experiments; S.Z.C. performed experiments; S.Z.C. and T.D.P. analyzed data; and S.Z.C. and T.D.P. wrote the paper.

## Supplemental materials

**Table S1.** Statistics for data collection, map reconstruction and model refinement

|  | Mg-ADP-P<sub>i</sub>-pyrenyl actin filament<br>(EMD-XXXXX, PDB XXXX) | Mg-ADP-pyrenyl actin filament<br>(EMD-XXXXX, PDB XXXX) |
| --- | --- | --- |
| **Data collection and map reconstruction** |  |  |
| Magnification | 47619 | 47619 |
| Voltage (kV) | 300 | 300 |
| Electron exposure (e<sup>-</sup>/Å<sup>2</sup>) | 55.87 | 55.87 |
| Defocus range (μm) (set/measured) | [-2.5, -1.5]/[-3.2, -1.3] | [-2.5, -1.5]/[-3.5, -1.3] |
| Pixel size (Å) | 1.045 | 1.045 |
| Rise (Å)/twist (°)/Symmetry | 27.37/-166.57/C1 | 27.39/-166.58/C1 |
| Initial particle images (no.) | 462669 | 268618 |
| Final particle images (no.) | 411301 | 240254 |
| Map resolution (Å) | 3.0 | 3.2 |
| FSC threshold | 0.143 | 0.143 |
| Map resolution range (Å) | 2.7-3.3 | 2.7-3.6 |
| Map sharpening B factor (Å<sup>2</sup>) | -100.43 | -114.39 |
| **Model Refinement** |  |  |
| Initial model used (PDB ID) | 6DJN | 6DJN |
| Model resolution (Å) | 3.07 | 3.27 |
| FSC threshold | 0.5 | 0.5 |
| **Model composition** |  |  |
| Heavy atoms | 11848 | 11828 |
| Protein residues | 1488 | 1488 |
| Ligands (ADP/P<sub>i</sub>/Mg<sup>2+</sup>/pyrene) | 4/4/4/4 | 4/0/4/4 |
| **B factors (Å<sup>2</sup>)** |  |  |
| Protein | 75.51 | 74.31 |
| Ligands | 70.84 | 70.21 |
| **R.m.s. deviations** |  |  |
| Bond lengths (Å) | 0.007 | 0.013 |
| Bond angles (°) | 0.788 | 0.953 |
| **Geometry validation** |  |  |
| MolProbity score | 1.38 | 1.45 |
| Clashscore | 5.12 | 3.46 |
| Poor rotamers (%) | 0.00 | 0.00 |
| **Ramachandran plot** |  |  |
| Favored (%) | 97.48 | 95.50 |
| Allowed (%) | 2.52 | 4.50 |
| Disallowed (%) | 0.00 | 0.00 |

**Table S2.** Comparison of filament rise and twist in the final models and each filament particle.

| Sample | Rise in final model | Rise in each particle |
| --- | --- | --- |
|  | Twist in final model | Twist in each particle |
| ADP-P<sub>i</sub>-pyrenyl-actin filaments | 27.380 ± 0.001 | 27.111 ± 0.735 |
|  | -166.579 ± 0.001 | -166.641 ± 0.819 |
| ADP-P<sub>i</sub>-actin filaments | 27.329 ± 0.002 | 27.377 ± 0.389 |
|  | -166.528 ± 0.003 | -166.637 ± 0.619 |
| ADP-pyrenyl-actin filaments | 27.396 ± 0.001 | 27.234 ± 0.822 |
|  | -166.596 ± 0.001 | -166.652 ± 0.836 |
| ADP-actin filaments | 27.518 ± 0.002 | 27.388 ± 0.479 |
|  | -166.625 ± 0.001 | -166.607 ± 0.705 |

Note: Values are in the format of mean ± standard deviation.

**Figure S1.**
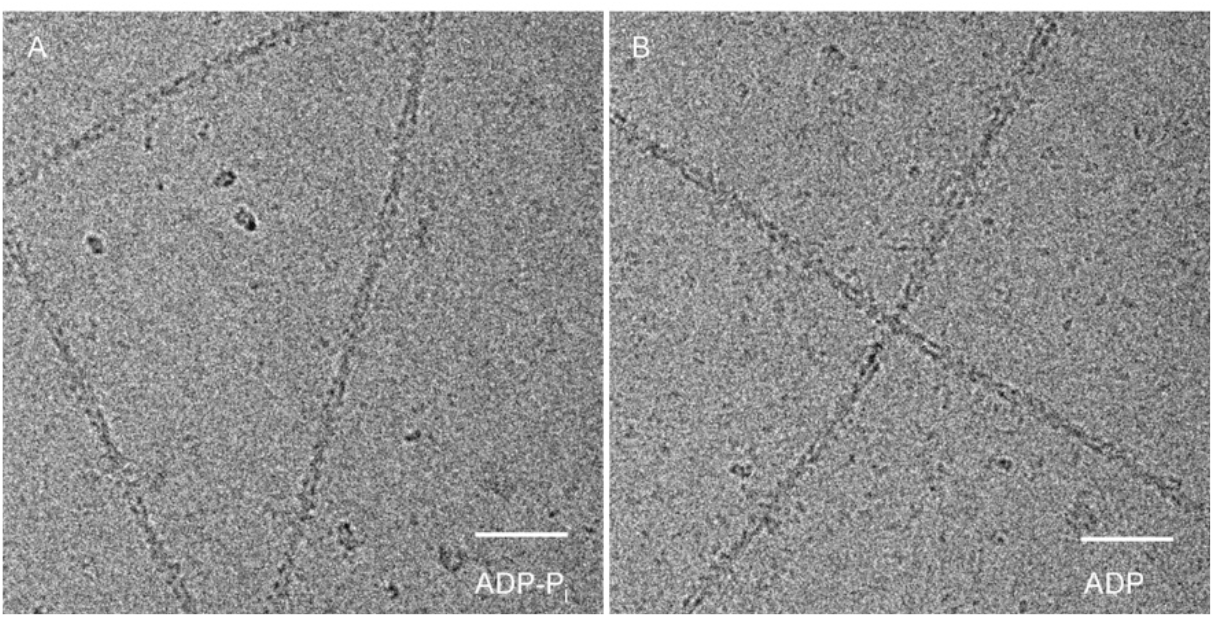
Micrographs of vitrified pyrenyl-actin filaments. (A) A representative area of ADP-P_i_-pyrenyl-actin micrograph. (B) A representative area of ADP-pyrenyl-actin micrograph. ADP-pyrenyl-actin has more unpolymerized monomers. Scale bar: 50 nm.

**Figure S2.**
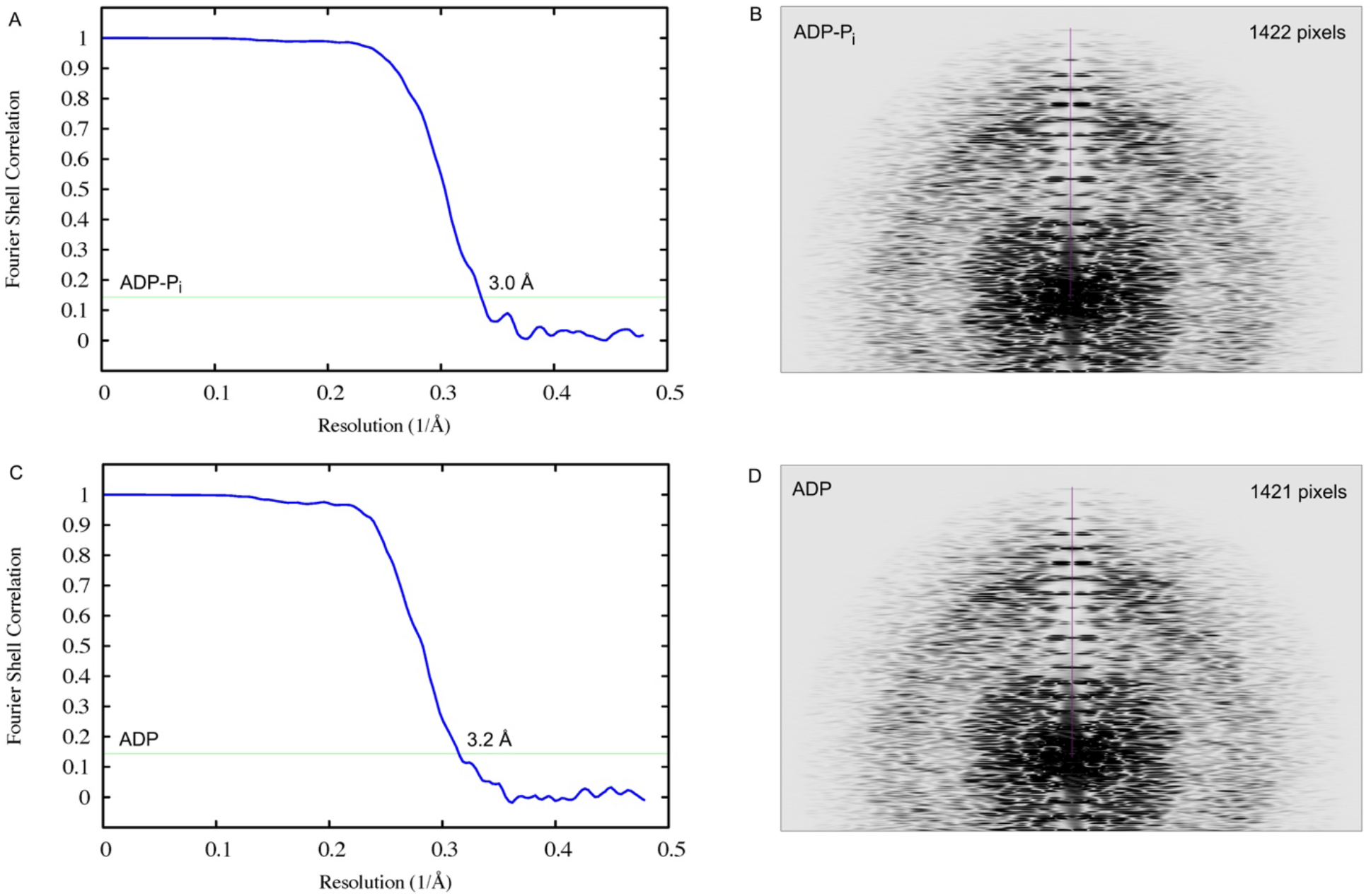
Global map resolution estimation. (A, C) Resolution of (A) ADP-P_i_-and (C) ADP-pyrenyl-actin maps estimated using Fourier shell correlation (FSC, blue curves) with 0.143 criterion (blue horizontal lines). (B, D) Resolution of (B) ADP-P_i_- and (D) ADP-pyrenyl-actin maps estimated using layer-line images calculated from map projections. The purple line is from the center of the layer-line image to the highest visible layer line. The size of layer-line images is 4096*×*4096 pixels, and the pixel size is 1.045 Å. Using the formula, resolution = (pixel size)*×*(layer-line image size)/(layer-line height), the estimated resolution is 3.0 Å for both ADP-P_i_- and ADP-pyrenyl-actin filaments.

**Figure S3.**
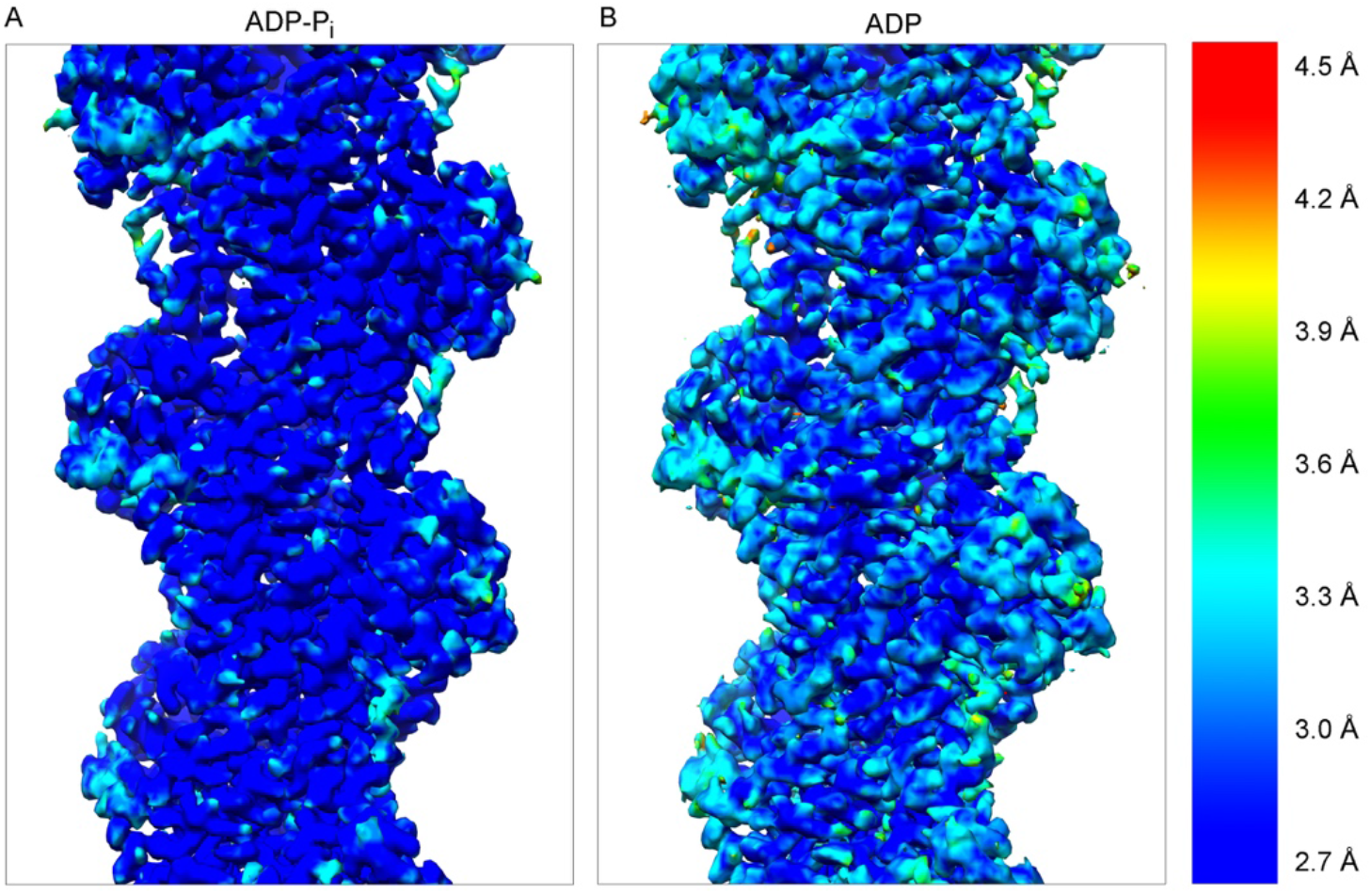
Local resolution estimation with ResMap. (A) Local resolution of ADP-P_i_-pyrenyl-actin filament. Resolution mostly falls in the range of 2.7 Å and 3.3 Å. (B) Local resolution of ADP-pyrenyl-actin filament. Resolution mostly falls in the range of 2.7 Å and 3.6 Å.

**Figure S4.**
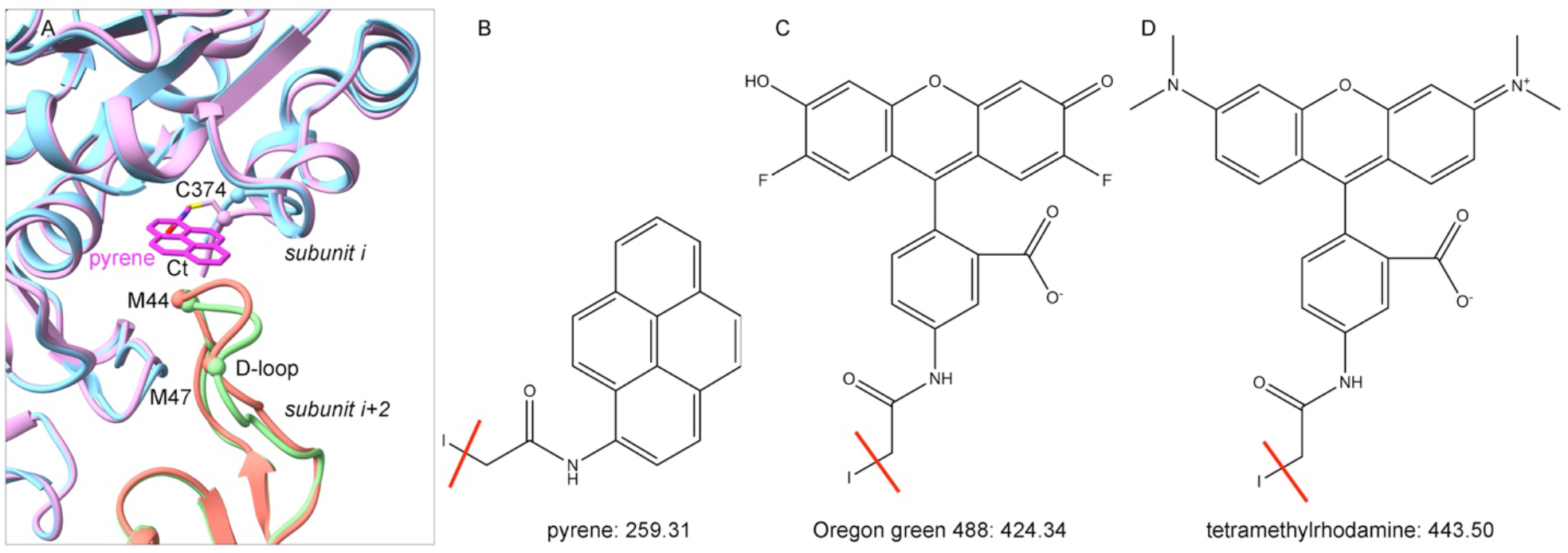
Pyrene binding site and structures of the commonly used fluorophores for labeling actin. (A) Conjugation of pyrene to C374 displaces residues between M44 and M47 in the D-loop. The two subunits (*i* and *i+2*) along the long-pitch helix in Mg-ADP-P_i_-actin filament are colored in sky blue and salmon, and subunits in Mg-ADP-P_i_-pyrenyl-actin filament are colored plum and light green. The pyrene group is colored in magenta. The C_α_ atoms of C374, M44 and M47 are shown as balls. (B) N-(1-pyrene)iodoacetamide (no charge). (C) Oregon green 488 iodoacetamide (a negative charge). (D) Tetramethylrhodamine iodoacetamide (a negative charge and a positive charge). During the conjugation reaction, iodine (on the left side of the red line) is removed. The molecular weight after the reaction is indicated below each structure.

